# Can transcranial electric stimulation with multiple electrodes reach deep targets?

**DOI:** 10.1101/382598

**Authors:** Yu Huang, Lucas C. Parra

## Abstract

To reach a deep target in the brain with transcranial electric stimulation (TES), currents have to pass also through the cortical surface. Thus, it is generally thought that TES cannot achieve focal deep brain stimulation. Recent efforts with interfering waveforms and pulsed stimulation have argued that one can achieve deeper or more intense stimulation in the brain. Here we argue that conventional transcranial stimulation with multiple current sources is just as effective as these new approaches. The conventional multi-electrode approach can be numerically optimized to maximize intensity or focality at a desired target location. Using such optimal electrode configurations we find in a detailed and realistic head model that deep targets may in fact be strongly stimulated, with cerebro-spinal fluid guiding currents deep into the brain.

Highlights
- Deep targets can be reached with intensities comparable to the cortical surface.
- Multi-electrode montages increase intensity with the same current limits per electrode.
- High-definition and intersectional pulsed stimulation are largely equivalent.
- Interferential stimulation is generally weaker than conventional stimulation.

## 1. Introduction

Conventional transcranial electric stimulation (TES) typically uses two sponge electrodes to deliver current to the scalp. Modeling studies suggest that electric fields generated with such an approach are diffuse in the brain and typically drop off in magnitude quickly with depth (Datta et al., 2009). A number of investigators have advocated the use of several small “high-definition” electrodes (Minhas et al., 2010) to achieve more focal or more intense stimulation (Datta et al., 2009; Dmochowski et al., 2011). This multi-electrode approach is sometimes referred to as high-definition transcranial electric stimulation (HD-TES) (Edwards et al., 2013; Dmochowski et al., 2013), and can be combined with any desired stimulation waveform. More recent efforts propose to use multiple electrodes with specific waveforms to reach deep targets in the brain (Grossman et al., 2017), or to increase intensity of stimulation (Vöröslakos et al., 2018). (Grossman et al. (2017) propose “interferential stimulation” with sinusoidal waveforms of similar frequency delivered through two electrode pairs. The investigators suggest that interference of these two waveforms results in maximal modulation of oscillating fields deep inside the brain. Vöröslakos et al. (2018) introduced “inter-sectional pulsed” stimulation, which distributes currents through multiple electrodes by temporal multiplexing, i.e. rapidly switching currents between multiple electrode pairs. This spreads out current on the scalp and minimizes skin sensation. The promise of non-invasive intense and deep brain stimulation has caused considerable excitement in the research community and popular media (Dmochowski and Bikson, 2017; Shen, 2018). For brevity, we will refer to these approaches as high-definition, interferential and pulsed stimulation, respectively.

Here we compare these three approaches in terms of the intensity and focality they achieve on the brain. We argue that high-definition and pulsed stimulation are essentially the same for the purpose of targeting. In contrast, interferential stimulation is genuinely different, but, as we will show, it achieves limited gains in depth of stimulation compared to the other two methods. The high-frequency waveforms of interferential and pulsed stimulation may have other advantages, such as a more robust electrophysiological response or reduced skin sensations, but for the purpose of targeting, as we will argue here, these three multi-electrode approaches are largely equivalent.

Having established this, we then address the central question of this paper, namely, can TES reach deep targets? We leverage numerical optimization methods developed for multi-electrode stimulation to achieve an optimal electrode montage on the scalp (Dmochowski et al., 2011). Optimization uses our latest detailed head model (Huang et al., 2016) with 0.5 mm resolution and computes optimal electrode montages for each location in the brain. This spatial survey shows that, generally, deep brain areas can only be diffusely stimulated. However, some preferred locations deep inside the brain can achieve field intensities that are comparable in magnitude to electric fields on the cortical surface. This is consistent with *in-vivo* recordings in human (Huang et al., 2017b), which show that fields in some deep brain areas can be as strong as in cortical locations. Thus, deep brain stimulation with TES appears feasible when using appropriate multielectrode montages, at least for some locations that are close to cerebro-spinal fluid (CSF).

## 2. Comparison of novel multi-electrode transcranial stimulation methods

We start with a review of three, seemingly distinct approaches of using multiple electrodes for transcranial electric stimulation. The emphasis will be on the intensity and distribution of the electric fields achieved by these multi-electrode methods.

### 2.1. High definition stimulation

Electric fields generated by more than two stimulation electrodes are additive, provided that current in each electrode pair is controlled independently (Dmochowski et al., 2011). Based on this observation, a number of investigators have proposed multi-electrode methods to maximize intensity or focality on a desired target in the brain (Dmochowski et al., 2013; Sadleir et al., 2012; Ruffini et al., 2014; Wagner et al., 2016; Guler et al., 2016; Dmochowski et al., 2017; Saturnino et al., 2017). The approach uses several small “high-definition” electrodes and can be combined with any arbitrary waveform. The term high-definition has been used most commonly for a specific electrode montage, know as “4×1” with a central electrode surrounded by four return electrodes (Datta et al., 2009, a better name would be “4+1” montage). This montage confines the area of stimulation with the 4 surrounding electrodes and thus achieves relatively focal stimulation. This was confirmed experimentally with supra-threshold stimulation in human (Edwards et al., 2013). On the flip side, the close proximity of the return electrodes leads to substantial shunting through the scalp and thus field intensities on the brain are weaker than with distant electrodes, and thus only reach cortical targets. Of course, this specific configuration of five electrodes is only one of many possible choices for arranging multiple electrodes. Here we will call high-defintion TES (HD-TES) any approach that delivers a given waveform simultaneously through multiple electrodes, regardless of the waveform used or the locations of the electrodes.

The additive property of current-controlled stimulation is particularly appealing for targeting as one can use efficient numerical optimization methods to find optimal electrode montages (Dmochowski et al., 2011). The approach is fairly flexible. One can optimize to achieving maximal intensity on a target, or one can attempt spatially constrained (focal) stimulation. Figure 1A shows an example of the electrode configuration that achieves optimal focal stimulation on a superficial target location (indicated with a black circle), while limiting the total current to 2 mA. The resulting montage has 6 electrodes and is shown on the left. It includes 1 anode (red) and 5 surrounding cathodes (green). This is a typical configurations when the desired field is to point in radial direction (Dmochowski et al., 2011). The field achieved at the target in this example is 0.5 V/m. To quantify how focal this stimulation is, we measure the brain volume that is exposed to an electric field of at least 0.25 V/m (i.e. half the magnitude at the target). In this specific example, this volume is 35 cm^3^, which corresponds to a length scale of 3.3 cm (cubic-root of 35 cm^3^). Note that current-flow often limits targeting in the sense that currents are higher on the path to the target, and so, as in this example, the maximal field is not necessarily at the target despite being optimally focal. In the second example (Figure 1B) we have maximized intensity on the target in radial direction and again limited the total current to 2 mA. In this case we selected a deep target. The resulting field intensity at this target location is estimated to be 1.7 V/m. The optimal electrode montage only uses two electrodes. Note that the electric field is also strong in areas that are distant from the target. Therefore, stimulation in this case is not focal. An advantage of the numerical optimization approach is that one can also constrain the current used in individual electrodes. For instance, if the total current is set to 2 mA, but the current through each electrode is limited to 1 mA, then the numerical optimization is forced to find optimal locations for at least 4 electrodes (at least 2 anodes and 2 cathodes, each conducting a maximum of 1 mA). Therefore, by limiting the current through each electrode, one can enforce currents to be distributed in space. Montages with 2+2 electrodes have been used previously in clinical trials with the goal of reducing skin sensations with small electrodes Dmochowski et al. (2013).

**Figure 1:**
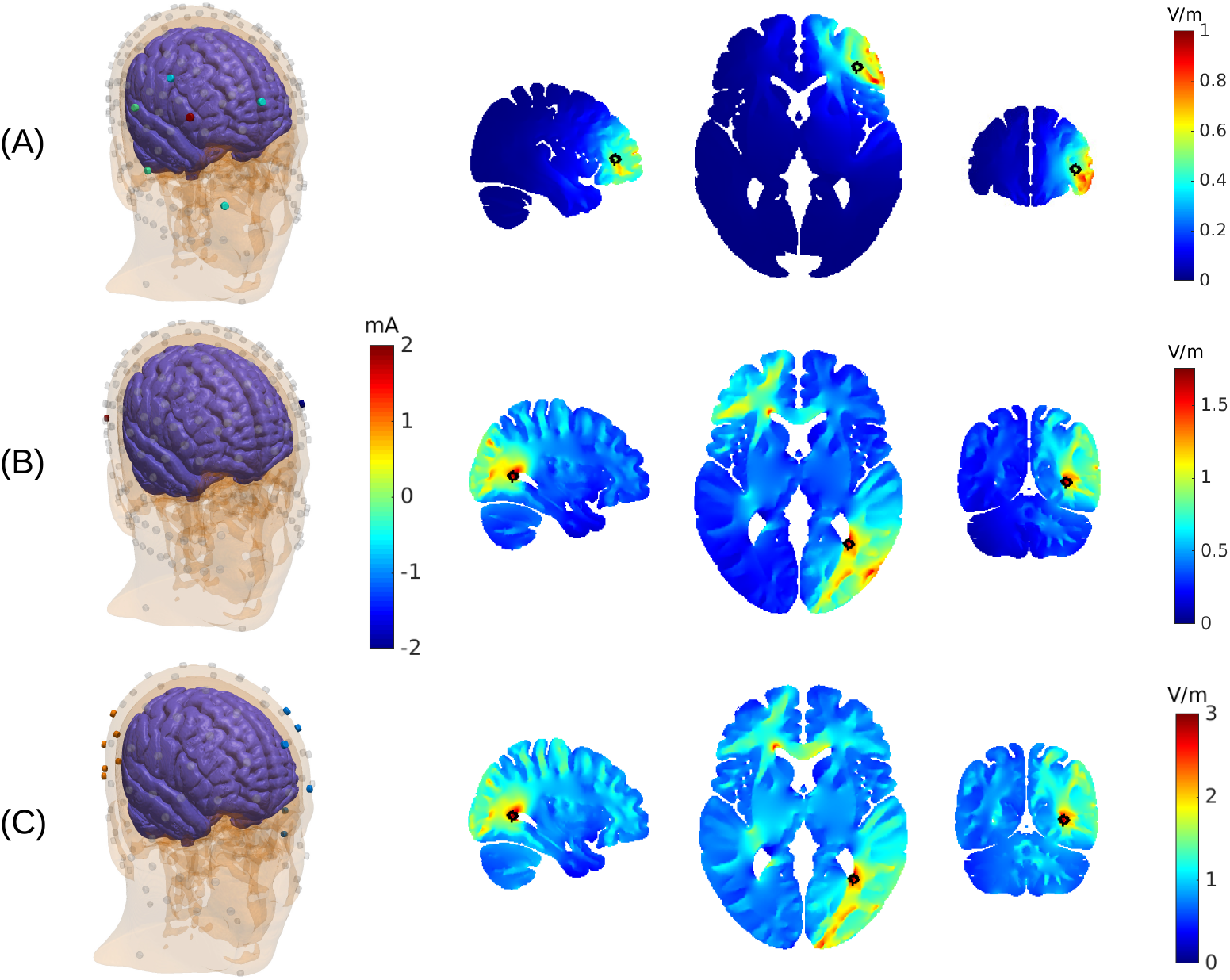
Examples of targeted stimulation using multiple electrodes. (A) Electrode locations and currents are optimized to achieve focal stimulation at a superficial target location (black circle). Total current is limited to 2 mA. Optimal locations are shown on the left with color indicating current magnitude through each electrode. Gray electrodes indicate possible locations considered by the algorithm, but which draw no current for this target. Resulting spatial distribution of electric field magnitudes in the brain is shown on the right. (B) Optimization here attempts maximally intense stimulation at a target location in the brain, again using 2 mA. The optimal solution only includes two small electrodes. (C) Here a total of 6 mA is used to target the same location as in (B), but current through each electrode is constrained to a maximum of 1 mA. Maximally intense stimulation is achieved by spreading the currents among 12 electrodes (6 anodes and 6 cathodes). This “high-definition” TES montage is equivalent to the “inter-sectional pulsed stimulation” protocol of Vöröslakos et al. (2018), but using optimal electrode locations.

### 2.2. Inter-sectional pulsed stimulation

Recently Vöröslakos et al. (2018) proposed a different approach to distribute currents in space, which they termed “inter-sectional pulsed stimulation” (ISP). Their idea is to rapidly switch currents between different electrode pairs. For instance, suppose that one wants to apply 6 mA through 6 pairs of electrodes (6 + 6 = 12 electrodes total). At any point in time the current is applied only to a single electrode pair, and each electrode pair conducts current only 1/6-th of the time. Thus, when averaged in time, each pair only applies an mean of 1 mA current. In one implementation, current switches every 60*μs* (a rate of 16.6 kHz). The basic argument of Vöröslakos et al. (2018) is that fields generated by each pair will polarize the cellular membrane of neurons in the brain, but because of the slow time-constant of the passive membrane (approximately 30 ms), polarization will be effectively low-pass filtered. The cell membrane blurs the rapid pulses in time, thus behaving effectively as it would in response to a continuous 1 mA current stimulation, which is the DC component of the monophasic 6 mA pulses. The amplitude of the pulses can be modulated at a lower rate and even reversed in sign. In this way, one can implement with ISP stimulation any desired waveform (that is slower than 1/30 ms=33 Hz). In Vöröslakos et al. (2018) this approach is demonstrated for 10 Hz sinusoidal stimulation equivalent to what is used in 10 Hz transcranial alternating current stimulation (tACS). Assuming a passive membrane (i.e. weak fields), the polarization of the membrane due to the different electrodes pairs will be additive (Weak fields of a few V/m do not trigger active current across the neuronal membrane, and thus it is fair to assume a passive membrane). This additivity includes the dependence of membrane polarization on field orientation, which will vary across electrode pairs. The motivation for the name “inter-sectional” is that this addition of fields will result in stronger fields wherever the individual field distribution of each electrode pair overlaps, or “intersects”.

As we discussed above, applying currents simultaneously also leads to an addition of the electric fields. Therefore, with the assumption of a relatively slow and passive membrane, the resulting effective field of ISP stimulation is identical to HD stimulation both in magnitude and orientation. Thus, for the purpose of spatial targeting, the ISP approach is numerically equivalent to the HD approach. In one case we assume temporal summation at the neuronal membrane, in the other case summation is strictly guaranteed by basic principles of electrostatics. Thus, for the purpose of targeting, we will treat the two approaches as one and the same. We can use the optimization approach presented above to maximize intensity on target while constraining the current through each electrode (Dmochowski et al., 2011). To follow the ISP scenario above, we allow for a total 6 mA while constraining the current in each electrode to no more than 1 mA. The optimization algorithm identifies 12 optimal electrode locations (6+6 montage) as shown in Figure 1C. The target location is identical to the example in panel B with a total of 2 mA. The field achieved at this target location is now 2.9 V/m. That is only about 2 times larger, despite the 3-fold increase in total current. On the other hand, this 2-fold increase in intensity is achieved with the same 1 mA maximum current at each electrode. If the limiting factor for sensation is the current at a single electrode location, then we have gained in intensity without an increase in sensation. Evidently, electrodes have to be sufficiently far apart in order to avoid cumulative effects on sensation of nearby electrodes (see Appendix A).

The approach of optimal targeting presented here is the same for HD and ISP. The methods, however, differ in one important aspect. With ISP stimulation currents are pulsing with a duty cycle of 1/6 at an intensity of 6 mA at each electrode location (for present example of a 6+6 montage). There is empirical evidence that this fast pulsing may lead to lower skin sensation as compared to the equivalent of a continuous 1 mA current (Paneri et al., 2016). Therefore, ISP stimulation may allow higher overall stimulation intensities by combining two distinct mechanisms. One is to distribute the currents in space (equivalent to using larger sponges in conventional TES, or spreading currents over many HD electrodes); and secondly, pulsing the currents at high frequency to reduce sensation on the skin.

### 2.3. Interferential stimulation with two pairs of electrodes

Another recent approach to using multiple electrodes for TES proposes the use of interfering AC waveforms (Grossman et al., 2017). The technique is used routinely in clinical practice in physical therapy, where it is known as “inteferential current therapy” (Goats, 1990). In the context of brain stimulation the idea was first proposed half a century ago (for a historical perspective see Guleyupoglu et al., 2013). Here we explore the ability interferential stimulation to reach deep brain areas (Grossman et al., 2017) and compare this to conventional HD-TES using the same set of electrodes.

The basic idea of interferential stimulation is that two sinusoidal waveforms of similar frequency, once added together, will generate a sinusoidal waveform that is modulated at a slower “beat” frequency (see Figure 2A). For instance, a sinusoidal wave of 2000 Hz added to a sinusoidal wave of 2010 Hz will be modulated in amplitude at 10 Hz. The strength of this modulation-the “modulation depth”-is defined as the difference of maximal and minimal amplitudes of the fast sinusoidal waveform, i.e. the range of the red envelope curve in Figure 2A. Of course, one can also generate a 2005 Hz waveform that is modulated in amplitude at 10 Hz and apply this stimulation waveform conventionally through the same set of electrodes. This is referred to here as “conventional” stimulation, although modulating sinusoidal waveforms is not currently common practice in TES. The interesting finding of Grossman et al. (2017) is that neuronal activity is entrained to this modulation frequency. In terms of targeting, the intuition of interferential stimulation is that this modulation with the slower beat frequency would be strongest, not at the surface of the scalp where the currents are applied, but in some depth in the brain where the two interfere. However, we note that the strongest intensity of stimulation is at the surface and thus the modulation depth is also strongest at the surface of the scalp (see Figure 2). Indeed, in Appendix B we show mathematically that the modulation depth of the interfering sinusoidal waveform is equal or smaller than what can be achieved with conventional TES with the same two pairs of electrodes. This is illustrated in Figure 2A with the modulation of amplitude shown as red and black curves for interferential and conventional stimulation, respectively. The result holds regardless of which field orientation one chooses to analyze (different columns in the figure represent different field orientations). Simulations on a realistic human head model (Figure 2) compare the field distributions achieved with two different electrode montages (analogous to simulations provided by Grossman et al. (2017) on a sphere or cylinder). The results for interferential stimulation do not fundamentally differ from conventional stimulation, except for magnitude, which is smaller with interferential stimulation (see histograms across the entire brain as “violin plots” in the third row of Figure 2B& C). Stimulation does appear to be more focal than conventional stimulation in some deep areas, but this result depends on the particular field orientations tested (Figure B.4). Details of the simulation and more detailed comparison between interferential and conventional stimulation are discussed in Appendix C.

**Figure 2:**
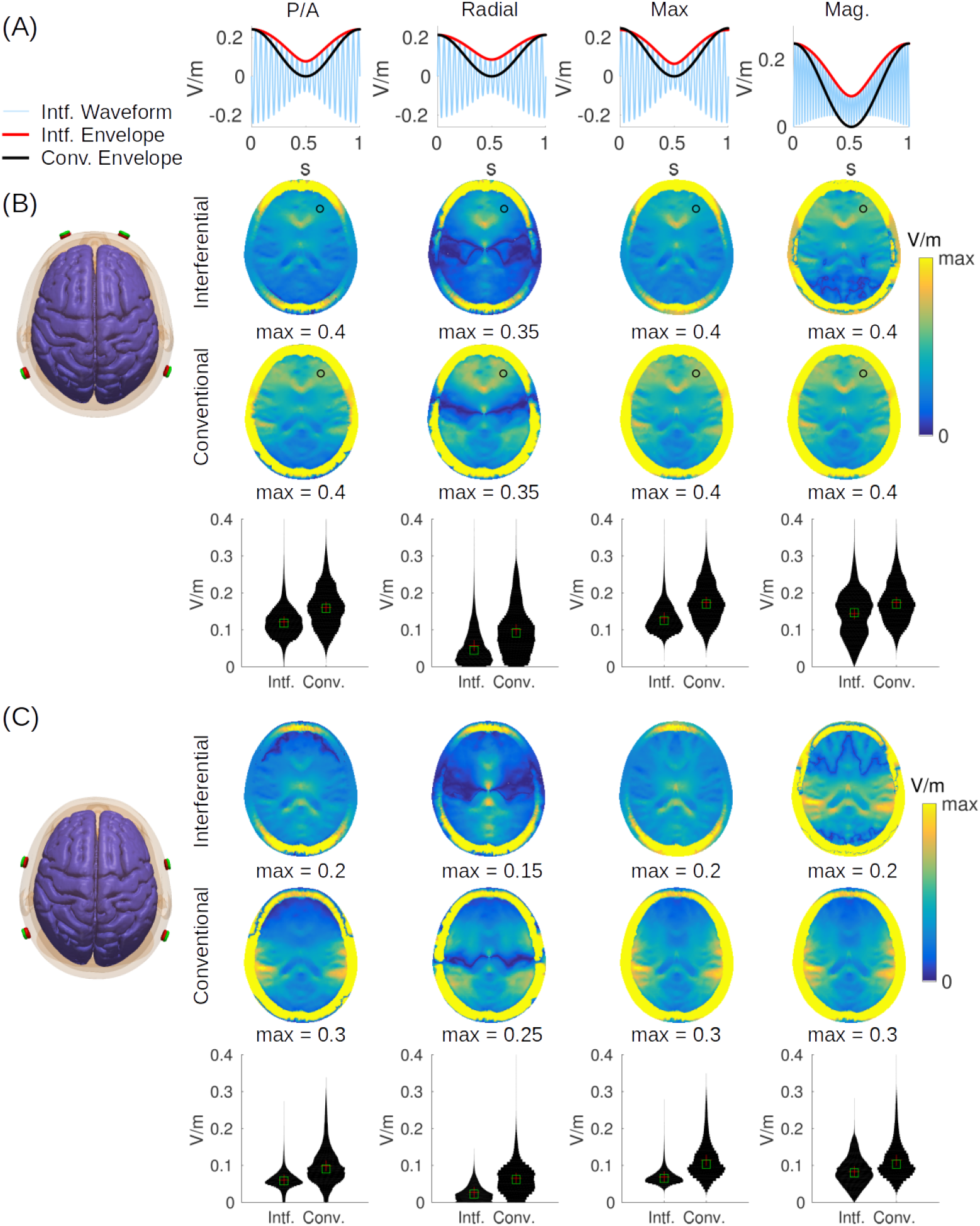
Modulation depth for interferential and conventional transcranial electrical stimulation. Stimulation is applied through two pairs of electrodes as shown on the left. (A) Waveform of interferential stimulation (light blue) along with the envelope for interferential (red) and conventional stimulation (black). Here we used 20 Hz as carrier frequency and 1 Hz as modulation frequency for better visualization. Waveforms were computed with values from the electric fields generated by each electrode pair (*a_i_* and *b_i_* in the equations of Appendix D) at the frontal brain location marked with a black circle in panel B. (B) In this configuration current flows between electrodes Fp1 and P7 on the left, and between Fp2 and P8 on the right, such that current flows predominantly in the posterior/anterior direction. (C) Here electrodes are closer together (with electrode pairs FT7/P7 and FT8/P8) generating more focal but also weaker stimulation. In panels B& C the first two rows show axial view of field modulation depth across the head (including skull and scalp). The third row indicates histograms across the entire brain. The four columns show the modulation depth of the electric field (*M*|_s_|, Eq. E.1) in three different directions (**n**): posterior/anterior (P/A), radial direction pointing to the MNI origin, i.e. anterior commissure (Radial), and the direction with the maximal modulation depth (Max). The last column is the modulation for the field magnitude (Mag.), or more specifically, the square-root of power modulation (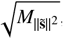Eq. E.5).

To summarize, in terms of field distribution, the three multi-electrode approaches provide similar results. Indeed, pulsed and high-definition stimulation can be regarded as equivalent, while interferential stimulation differs, but achieves similar results, with somewhat lower modulation depth. Furthermore, interferential stimulation is limited to producing amplitude-modulated waveforms, whereas pulsed and high-definition stimulation can be combined with arbitrary waveforms.

## 3. Is deep TES possible?

We now turn our attention to the central question of this paper, namely, can multi-electrode approaches be used to reach deep targets in the brain? In Figure 2 we have shown field distributions for two ad-hoc electrode montages, which are by no means optimal in terms of reaching deep targets. Perhaps by optimizing stimulation montages, as demonstrated in Figure 1, one can in fact reach various deep targets. In Figure 1 we tested two particular locations in the brain, one more superficial and one deeper (not coincidentally, we selected a location close to a ventricle). To determine whether these results are typical for other superficial or deeper locations in the brain we repeat the process surveying the entire brain. For each brain location we optimize the electrode montage to achieve either maximal focality or intensity, while limiting the total current to 2 mA. These alternative optimization criteria are important because there is a direct trade off between intensity and focality of stimulation, and one cannot achieve both simultaneously (Dmochowski et al., 2012). Figure 3A shows the size of the stimulated volume when optimizing for focal stimulation of each brain location. The volume encompasses all brain areas for which the field magnitudes are at least 50% of the field at the target, regardless of where in the brain this is (false-color in this figure indicates size on a centimeter scale as the cubic-root of this volume). This figure confirms our expectation that focal stimulation of deep brain areas is difficult, whereas on the cortical surface stimulation can be focal. The reason for this is that in order to reach deep brain areas currents have to pass through more superficial areas.

**Figure 3:**
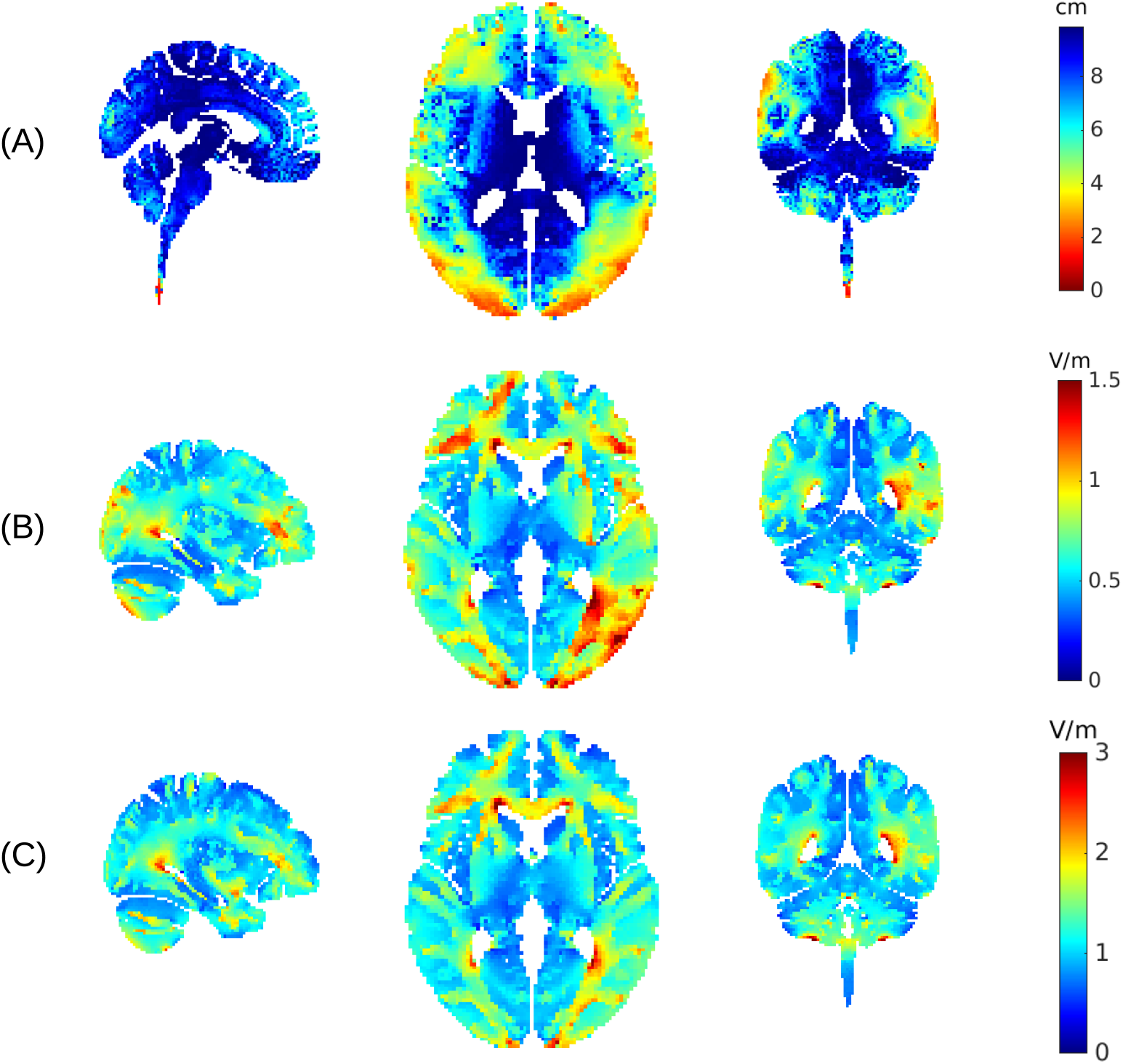
Distribution of achieved focality and intensity across the brain. (A) For each location a separate optimal electrode montage is found with the goal of achieving focal stimulation. The optimal montage is evaluated for the resulting size of the stimulated volume. Color indicates the size of the volume stimulated at 50% (or more) of the intensity at the target (we show the cubic root of the volume to indicate the length scale in centimeter). (B) Electrode montage is optimized, for each target location, to achieve maximal intensity with 2 mA. False color indicates the maximal value achieved at that location. (C) Same as B, but now injecting a total of 6 mA while constraining the current in each electrode to no more than 1 mA. Targeting was performed using a detailed 0.5 mm resolution current-flow model (as shown in Figure 1), but target locations were surveyed only on a 2 mm grid.

Figure 3B shows the maximal intensity that can be achieved at each location of the brain, when optimizing for intensity. Here it becomes clear that some deep brain areas can have stimulation intensities quite similar to the cortical surfaces. These tend to be located close to ventricles with highly conducting cerebro-spinal fluid (CSF), which serves as conduit to these locations. We also optimized for intensity, but now increasing total current to 6 mA while limiting current through each electrode to 1 mA (Figure 3C). This choice is again motivated by the desire to increase stimulation intensity in the brain, while limiting sensation on the scalp. The results are nearly identical to Figure 3 B, in terms of depth of stimulation, but now evidently with stronger field magnitudes reaching values of up to 3 V/m.

An ancillary empirical finding of these exercises is that maximizing for intensity always results in the smallest possible number of electrodes, given the current constraints. Limiting currents to a maximum value at each electrode (and in the sum) results in a set of linear constraints (Dmochowski et al., 2011), which apparently strictly favours sparse solutions. For example, with a total of 2 mA and 2 mA allowed through each electrode, the algorithm never chooses to distribute beyond a single pair. Similarly, for 6 mA total, with a 1 mA constraint per electrode, the algorithm always selects 6 cathodes and 6 anodes. In contrast, when optimizing for focal stimulation the algorithms chooses to distribute currents among more electrodes than the necessary minimum. Solutions range from 2 to 14 electrodes with a median of 7. For deeper targets the number tends to increase, consistent with the theoretical results on a spherical model (Dmochowski et al., 2012).

## 4. Discussion and conclusion

In summary, regardless of whether currents have to be distributed on the scalp or not, there are locations deep inside the brain that may be reached with fairly strong electric fields. The specific locations tend to be close to ventricles and deep sulci, which can guide currents along highly conducting CSF. Whether or not these computational predictions are accurate in terms of the exact locations of higher intensity stimulation has to be empirically validated. A particular caveat is that CSF in sulci and around the brain forms a very thin layer and therefore is hard to accurately model. We therefore have used the best available model to-date in terms of spatial resolutions. This model is based on the latest MNI dataset at 0.5 mm resolution and is particularly well suited for current-flow modeling as it extends the field-of-view to include the neck (Huang et al., 2016). Nevertheless, these model details, in particular sulci, are not reliably reproduced with lower resolution models. Location, depth and width of sulci also drastically vary from one subject to the next. Thus relying on gyral folding for the purpose of targeting will be associated with significant uncertainty across subjects. Additionally, tissue conductivity may not be isotropic or uniform and thus one should reserve some healthy skepticism on the detailed predictions of hotspots in computational models. However, targets located along the main ventricles and the central fissure may be more reliably targeted across subjects. The latest empirical evidence (Huang et al., 2017b; Chhatbar et al., 2018) supports the general take-home message of this work, namely, that deep brain areas may be stimulated with comparable magnitudes to cortical areas, at least if the electrodes are appropriately placed. Future modeling studies could survey specific brain areas to determine whether they can be consistently targeted across subjects.

While we have argued that high intensities may be achieved in depth, we also argue that such stimulation is not focal, because, to reach deep brain areas currents necessarily have to pass through superficial areas. To avoid accidental stimulation along the way one may use optimization algorithms that can minimize stimulation at undesirable locations (Huang et al., 2018).

Note that maximal electric fields reported here of about 3 V/m are quite a bit larger than the previous modeling literature, which generally estimates fields to be less than 1 V/m in the human brain when using 2 mA current on the scalp. This has multiple reasons. First, we used optimal electrode montages to maximize intensity; second, the model employed small electrodes which are known to generate stronger fields in the brain (Dmochowski et al., 2012); and third, when we do distribute currents among many electrodes, we also inject significantly higher total current than conventional sponge TES. Increasing field magnitudes by distributing currents in space was an important contribution of the work by Vöröslakos et al. (2018). However, note that despite a 3-fold increase in the total current injected, fields only increase by a factor of 2. This is because the currents have to be distributed on the scalp surface, and thus we lose some of the benefit of ideally placing small electrodes.

Optimization was based on the conventional linear superposition principle introduced in the context of high-definition TES (Dmochowski et al. 2011). We argued that this linear superposition applies identically to inter-sectional pulsed stimulation. Therefore, our results obtained here should apply to both methods. We were not able to identify scenarios for which interferential stimulation outperformed conventional stimulation. The conventional stimulation, which simply adds the fields of multiple electrodes pairs, achieves generally larger field magnitudes and modulation. The size of focal spots with interferential stimulation appears smaller in some directions, but this is not a consistent finding across all field orientations, at least for the electrode configurations tested here. Here we used montages resembling those proposed in the original work of (Grossman et al. 2017), but there may be other montages where the benefits of interferential stimulation are more apparent. At present the optimization of conventional high-definition TES seems to achieve more focal stimulation for superficial targets and more intense stimulation in depth. Nevertheless, we cannot rule out that more sophisticated non-linear multivariate optimization of electrode location for interferential stimulation could change something to this picture (e.g. Cao and Grover, 2017).

Both inter-sectional pulsed stimulation and interferential stimulation use high-frequency waveforms. However, inter-sectional stimulation used monophasic pulses, which, when low-pass filtered by the cellular membrane, behave effectively as slow (or DC) stimulation. In contrast, interferential stimulation uses biphasic sinusoidal wavforms, which, when low-pass filtered, should give no net membrane polarization. The unexpected empirical observation of (Grossman et al. 2017) is that neural activity entrains to the slower amplitude fluctuations of the modulated sinusoids, which cannot be explained with a conventional passive membrane response.

In all simulations we have assumed a quasi-static solution to Maxwell’s equations. Current induction in tissue with weak fields (~ 1 V/m) and signal propagation at low frequencies (~ 10 kHz) can be rightfully neglected (Plonsey and Heppner, 1967). Capacitive effects at low frequencies (< 1 kHz) may be mostly attributed to the electrodes-tissue interface (Schwan and Kay, 1957), while capacitive effect of tissue based on dielectric constant measurements (Gabriel et al., 1996b) should be negligible (< 1 degree, see Supplementary Figure S3). However, tissue conductance values do change with frequency (Gabriel et al., 1996b) and thus we reproduced the present figures with conductivity values cited in the literature for 1 kHz (Gabriel et al., 1996a,1996b; Baumann et al., 1997). The results are qualitatively the same to those shown in the present Figures 2 and Figures B.4 (see Supplementary Figure S4 and Figure S5).

**Figure 4:**
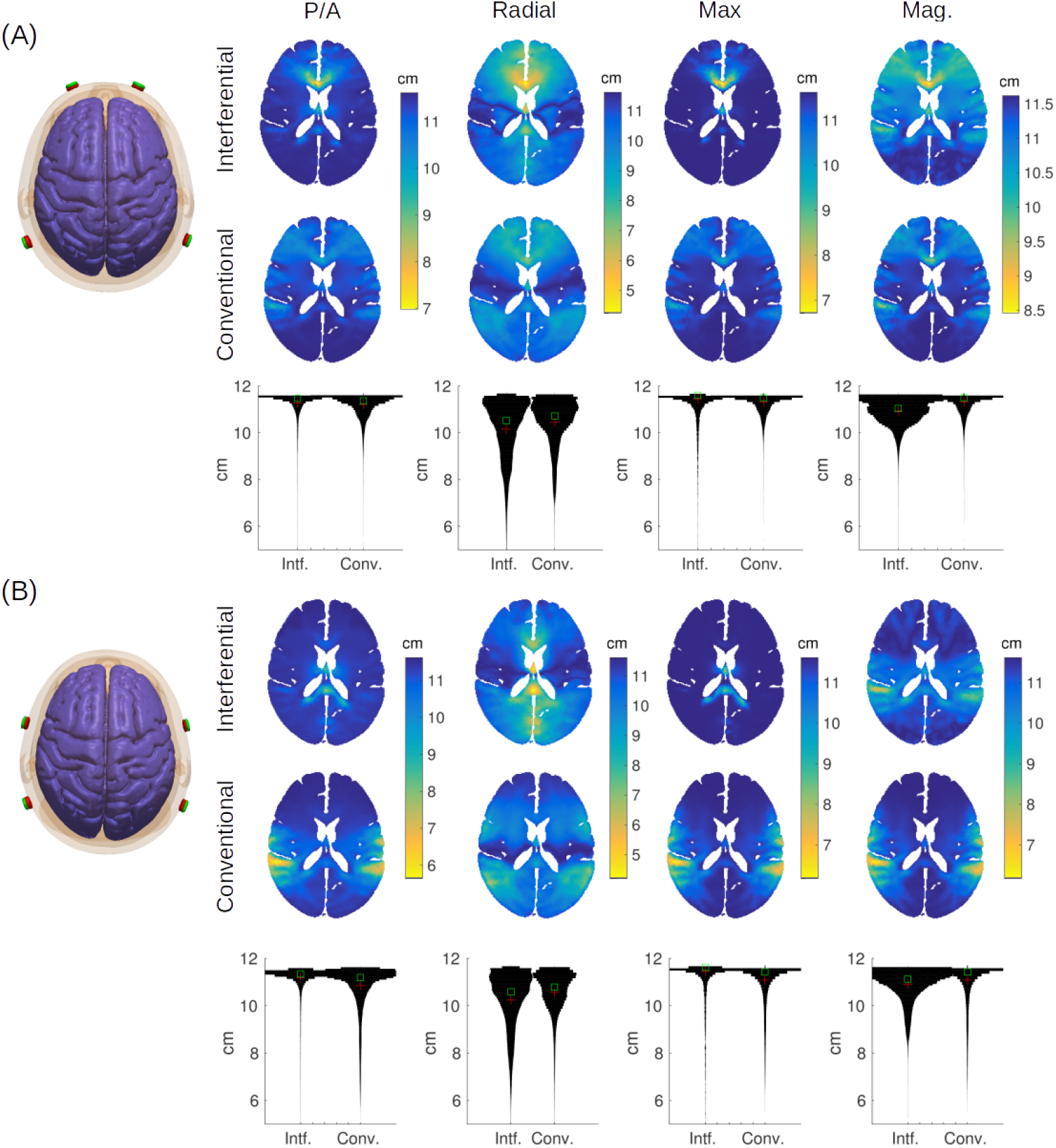
Figure B.4: Focality for interferential and conventional transcranial electrical stimulation. Same simulations as shown in Figure 2 except here spatial extent of stimulation is evaluated for each location. Color indicates the size of the volume stimulated at 50% (or more) of the intensity found at each location (again, showing the cubic root of the volume to indicate the length scale in centimeter).

It is important to note that in all cases optimization attempted to achieve radial fields at the target location. Differing field orientation may result in somewhat different preferred locations for targeting. We expect, however, that targets close to CSF can be preferentially reached regardless of orientation.

## 5. Acknowledgements

We would like to thank Gyorgy Buzsaki, Jacek Dmochowski and Marom Bikson for their comments on an earlier version of this manuscript. We also thank Gavin Hsu for his help on the recordings for Figure S3. This work was supported by NIH grants R01MH111439, R01MH111896 and R44NS092144.

## Appendix A. Targeting methods

For targeting we uses an existing detailed head model including neck with 0.5 mm resolution and 231 electrodes (Huang et al., 2016, see the 3D rendering of the head in Figure 1A). Briefly, each voxel of the model is categorized into different tissues with corresponding uniform tissue conductivity (scalp, skull, CSF, gray and white matter, and air cavities). Electrodes are placed according to the 10/05 system with two additional lower rows of electrodes as well as 4 electrodes on the neck to allow more flexibility in current flow. The neck electrodes, in particular, serve as a reasonable approximation of extra-cephalic electrodes often placed on the torso or arms. The model is then meshed into a volumetric tetrahedral mesh (using ScanIP) and electric field distributions estimated using finite-element modeling of current flow (using Abaqus). The resulting field estimates for each electrode (relative to a common reference) are then resampled in the original volume of 0.5mm resolution (for details on this entire modeling process see Huang et al., 2016).

Targeting largely follows (Dmochowski et al. 2011). We scanned the brain at 2 mm resolution leading to a targeting survey on about 210k brain voxels including gray and white matter (scanning at the full 0.5 mm resolution was computationally prohibitive). For focal stimulation we selected the LCMV approach with an L1-norm constraint. The L1-norm constraint results in sparse solutions, i.e. solutions use a small number of electrodes. In all cases we have optimized in radial direction, that is, we have maximized for field magnitude in the direction pointing towards the center of the brain. For convenience we have selected the origin of the MNI coordinate system, which places the anterior commissure as its center (Evans et al., 1993). It is important to note that the detailed results do significantly depend on this choice. In particular, when we are close to CSF, the strongest fields are achieved in the direction orthogonal to the CSF/brain surface. This sometimes coincides with radial direction, but not always. We have nonetheless selected radial direction, since selecting preferred orientation is somewhat arbitrary in deep brain areas. An alternative approach would have been to search for the optimal orientation (Dmochowski et al., 2013), i.e. optimal stimulation regardless of orientation. Unfortunately this approach has no efficient numerical solutions. Thus, finding an optimal stimulation montage, irrespective of orientation, is computationally challenging for the large number of locations, which we optimize in this targeting survey across the brain.

When constraining the currents through individual electrodes and thus distributing them to reduce skin sensation, it is important to provide sufficient spacing between electrodes. Otherwise, close-by electrodes will result in significant summation of electric fields on the scalp. Therefore, for Figure 1C and Figure 3C we used a subset of the 231 electrodes, as shown in the 3D rendering in Figure 1C.

## Appendix B. Modulation depth with interferential stimulation

Electric current applied to the surface of the head generates electric fields of varying intensity throughout the tissue. When multiple current sources are applied, the electric field vectors add together. Let’s assume we have two separate pairs of electrodes, and they achieve at a particular location in the tissue a field with amplitudes *a* and *b* respectively, say, in anterior/posterior direction. If each electrode pair applies an alternating current with sinusoidal waveforms of different frequencies, then the addition of these fields will interfere and result in a modulated sinusoidal waveform (Figure 2A). An alternative approach, which we refer to as the “conventional” stimulation, simply applies the same *modulated* sinusoidal waveforms to both electrode pairs. The resulting envelope curve is shown in black (Figure 2A). By comparing the red and black envelopes, it is easy to see that the modulation depth cannot be any larger for interferential stimulation than for conventional stimulation. Indeed, basic algebra presented in Appendix D indicates that the modulation depth for interferential stimulation is 2 * min(a, b) (see Eq. D.4), whereas for conventional stimulation it is *a* + b, which is larger except where *a* = *b* (We are assuming *a* and *b* are positive, but the sign of stimulation can always be flipped if they are not). Thus, conventional modulated transcranial AC stimulation (modulated tACS) can achieve the same or stronger modulation depth with the same peak current intensities applied to each electrode pair. What about direct current stimulation (tDCS)? In that case field intensity is also simply *a* + *b*. So conventional TES can achieve the same or stronger field intensity (with tDCS) or modulation (with tACS) than interferential stimulation.

This result holds regardless of which field orientation one considers. But perhaps the result is different when we consider the *magnitude* of the electric field instead. After all, (Grossman et al. 2017) found that the neuronal response is synchronous with the lower beat (modulation) frequency of 10 Hz, regardless of the reversing polarities, which oscillate at 2005 Hz. If all that matters is the strength of the field, regardless of orientation, then field magnitude may be the better metric. So how is magnitude (or equivalently power) of the oscillating field modulated in the case of interfering stimulation? The power is modulated in complicated ways (Figure 2A, 4th panel), however, at the relevant beat frequency, one can calculate the modulation depth explicitly (Eq. E.5). Here, again, we find that the conventional approach achieves the same if not stronger modulation depth in the magnitude (power) of the field vector, provided the sign of stimulation is appropriately selected (see Appendix E). Again, this can be seen readily without any math, by comparing the red and black curves (Figure 2A, 4th panel).

These results are entirely independent of where in the head we are measuring modulation depth or how strong the fields can be at that location. Instead, they are a basic mathematical property of interfering sinusoidal waveforms as we derive in Appendix E.

## Appendix C. Focality of interferential and conventional stimulation

While the conventional approach discussed in the previous section may achieve stronger modulation at a given location, it is not clear whether this stimulation is stronger at all brain locations or whether it is more focal. Perhaps with interferential stimulation, fields cancel fortuitously in some locations and add up constructively in others, so as to achieve more focal stimulation in deeper brain areas. The specific details of current flow may be important in this context. We therefore simulated field distributions for the inhomogeneous anatomy of the head, which has a complex 3D morphology involving skin, skull, highly conducting CSF, and a gyrated brain. These simulations were based on a finite element model of a standard human head from the Montreal Neurological Institute with 1 mm resolution (MNI-152, Grabner et al., 2006). Electric field distribution with the two electrode pairs placed at locations Fp1/P7 and Fp2/P8 were simulated using ROAST, an easy-to-use toolbox for realistic current-flow models of the human head (Huang et al., 2017a). With this, we generated the distribution of the electric field intensity for each electrode pair and direction (These signed intensity values are the *a_i_* and *b¡,* for 3 spatial directions *i*=1… 3 used in Appendix E). From these values one can compute the modulation depth of the field in any arbitrary direction as well as the modulation depth of the field power (Eqs. E.1 and E.5). The resulting modulation depth for the interferential stimulation are generally smaller than for the conventional approach (Figure 2B& C), regardless of field orientation. This is particularly evident in the histograms that compare modulation depth across the entire brain (third row of panels B& C). The results do not change when we bring electrodes closer together (comparing Figure 2B with Figure 2C).

To determine whether the stimulation is more focal, we evaluate in Figure B.4 the extent of the stimulated volume for the same montages and field orientations as in Figure 2. This is quantified as the brain volume that has a field of at least 50% of the intensity at the current location. Size of this volume is shown as false-color images, i.e. color and not the size of a hotpot is indicative of the stimulated volume. For radial direction, interferential stimulation has mid-line locations with more focal stimulation volume as compared to the conventional approach.

Histograms of the size of stimulated volume across the entire brain (“violin” plots in the third row of panels A& B) indicated that the vast majority of voxels are not focally stimulated (maximum value of 11.62 cm corresponds to the entire brain). But there are a small fraction of voxels that have a more narrow focus (values below 6 cm). Radial direction and overall magnitude accentuate this, and for these there is an advantage for interferential stimulation. However, this is not necessarily true for the other field orientations (P/A, Max), which appear to be more focal with the conventional approach, at least for superficial areas.

In these simulations focal spots appear in the midline because current strengths are equal on either side of the brain. We also simulated currents in a 4/1 ratio between left/right electrode pairs with the total injected current held at 2 mA (following the example of Grossman et al., 2017). As expected, the locations of strongest modulation move laterally (see Supplementary Figure S1). Also, hotspots are somewhat less well defined because midline CSF no longer aid current flow to midline locations. Additionally, the benefit of conventional stimulation in terms of modulation depth becomes stronger because for interferential stimulation modulation depth is weaker when field intensities are unequal (see Appendix D& Appendix E). In terms of focality, the results (Supplementary Figure S2) are comparable to what is shown in Figure B.4.

## Appendix D. Modulation envelope in 1D

Here we derive equations for the depth of modulation when two sinusoidal waveforms are added together.

Take two sinusoidal waveforms with frequencies *ω_a_* and *ω_b_* and add them together with amplitudes *a* and *b*:

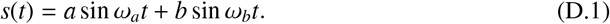

This combined waveform will “beat”, which means that a fast oscillation with frequency 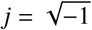 is modulated in amplitude with the “beat frequency”, *Δω*=*ω_a_ - ω_b_* (light blue curve in Figure 2A). The extent of this amplitude modulation depends on the individual amplitudes *a* and *b*. To determine the modulation “depth” of this beating signal, it is helpful to compute the envelope (red curve in Figure 2A), which can be defined as the absolute value of the analytic signal:

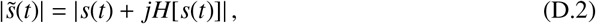

where H[·] stands for the Hilbert transform, 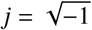and |·| is the absolute value of the complex numbers. Plugging Eq. D.1 into Eq. D.2, noticing that the Hilbert transform of a sin *ωt* is simply a cos ω*t* we get 

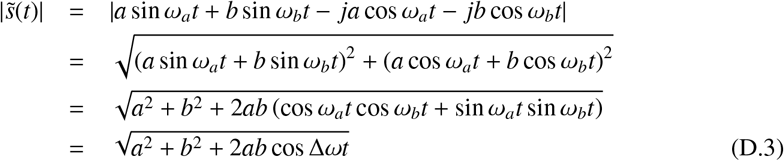

Here we assumed, *ω_a_> ω_b_,* so that the beat frequency is positive. We define the modulation depth as the difference between the maximal and minimal value of the envelope. From Eq. D.3 we can simply determine the maximum and minimum of the envelope, as it only depends on a single cosine, cos Δωt, with maximum/minimum values of ±1. The modulation depth is therefore:

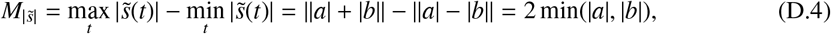

A conventional approach of stimulation simply applies current simultaneously (either DC or AC) and the total field amplitude is additive. This would be simply *a* + b. Of course this can be modulated say in the case of AC with a cosine as follows:

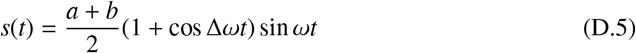

In this case the modulation depth is simply *|a* + *b*|, and since we are free to pick the polarities (in case they are not the same sign), that results in |a| + |b|. Evidently, |a| + |b| ≥ 2min(|a|, |b|).

For interferential stimulation using three or more sinusoids (e.g Cao and Grover, 2017), one can also readily show that |a| + |b| + |c| > M|_s_|, where *c* is the amplitude of the 3rd sinusoid. Therefore, a conventional TES approach can always be made to reach stronger modulation depth than interferetial stimulation.

The modulation index is defined as the modulation depth over the mean amplitude. For the interfering sinusoids this is:

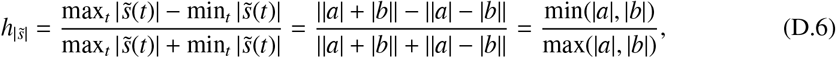

which has values between 0 and 1.

With the conventional approach the modulation index can be selected arbitrarily, regardless of values *a* and *b*. With the choice of modulation as in Eq. D.5, the modulation index is 1 everywhere.

## Appendix E. Modulation in 3D

Let’s now assume we have a 3D signal, *s*(*t*) = [s_1_(*t*), *s*_2_(*t*), *s*_3_(*t*)], that is the sum of two 3D sinusoids *s*_i_(*t*) = *a_i_* sin *ω_a_*t* + *b*_i_* sin *ω_b_t*, *i* = 1… 3. The modulation depth and index for the interferential stimulation obtained from Appendix D still hold along a specific direction represented by a unit vector **u**, i.e.,

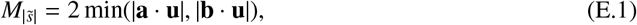

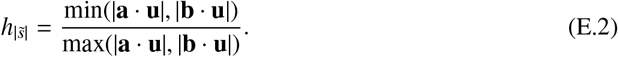

In particular, one may be interested in the direction along which the modulation is maximized. It is straightforward to show that Eq. E.1 is maximized if and only if **u** falls onto the plane spanned by **a** and **b**. Therefore, the maximal modulation depth can be obtained by

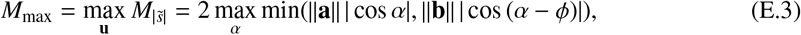

where *φ* is the angle spanned by **a** and **b**, and *α* is the angle between a and the projection of **u** on that plane. We do a 1D-search over *α* e [0,2*π*) to find the maximal modulation *M*_max_ for values ||**a**||, ||**b**||, *φ* in each location of the brain.

Note that for conventional stimulation (with modulation as in Eq. D.5) the modulation in direction **n** is simply (**a** + **b**) · **n** and the maximum ||**a** + **b**||. Because the proof provided for 1D above applies in any direction **n** we know again that interferential stimulation cannot be stronger than conventional stimulation, including for the direction with maximal interferential stimulation.

For the modulation of the magnitude of the 3D field, we define the following as the power envelope:

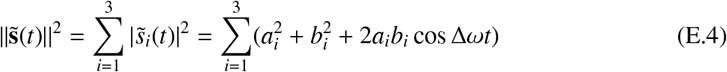

This quantity does not capture the envelope of the magnitudes ||**s**(*t*)||^2^=Σ*_i_* |s*i*(*t*)|^2^ of the 3D vector (light blue curve, Figure 2A, 4th panel). However, it does correctly capture the power modulation *at the beat frequency* Δω (red curve, Figure 2A, 4th panel). With a similar definition for modulation depth, but now for the power envelope at the beat frequency, we obtain

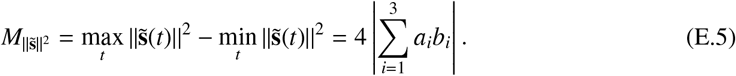

The modulation depth for the conventional case is simply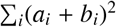. But we could have also flipped the sign of stimulation to achieve 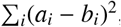, if that value is larger. One can show that 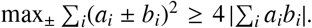. This means, once again, that modulation depth can always be made larger with the conventional approach, if the sign of stimulation is properly chosen.

The modulation index for the power is

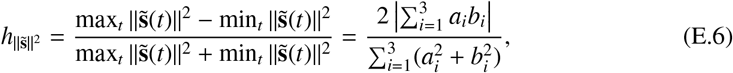

which is between 0 and 1.

## Supplementary Information: Can transcranial electric stimulation with multiple electrodes reach deep targets?

Here in this Supplementary Information, we address two questions related to the interferential stimulation: 1) how unequal input currents at the two electrode pairs can change the electric field distribution? and 2) is the quasi-static assumption of the underlying physics appropriate for stimulation in the kHz range?

### Interferential stimulation with unequal input currents

The results shown in Figure 2 and Figure B.4 simulate equal current on both electrode pairs. Below are the same figures for the case that current intensities on the two pairs have ratio of 4/1 (left vs. right pair, the sum is held at 2 mA). This is the ratio used in an example provided by (Grossman et al. 2017) in their Figure 2E. Our Figure S1 shows that, as expected, the areas of highest modulation move laterally. Interestingly, the focal spot moves in opposite directions for the two methods. A leftward move is expected for conventional stimulation as higher modulation will be closer to the electrodes with the highest current intensities. Modulation for interferential stimulation is maximal where fields are equal (in magnitude and direction). That area moves rightwards, as we have to be closer to the weaker electrode for the two fields to be the same. We also see that the modulation depth differs now more significantly between interferential and conventional (with modulation stronger for conventional, Appendix D& Appendix E)). When comparing Figure 2 and Figure S1 it also appears that the location of focal spots in depth relate mostly to anatomical details.

As with Figure B.4, Figure S2 shows that the size of the focal spot is smaller for interferential stimulation, but this is not consistent across all directions considered (again, note the volume of stimulation is represented here as false-color, while the size of the hotspots is not as meaningful).

### Quasi-static assumption is valid for kHz range stimulation

To be sure that capacitive effects can be neglected we measured phase delays in tissue (on the arm) *in vivo.* A single subject participated in this study after providing signed consent following the guidelines of a protocol approved by the IRB of the City University of New York. We applied a sinusoidal current with 0.5 mA peak-to-peak amplitude between two stimulation electrodes using a current-controlled stimulator, and measured the resulting voltages relative to a common reference at two intermediate locations on the arm separated by about 4 cm (see Figure S3A). In the range of 800 Hz-10 kHz we find a phase delay of less than 1 degree between the two locations (less than 1/360 of the sinusoidal cycle duration). This negligible delay confirmed expectations from prior literature and thus we did not measure additional subjects. Given that the dielectric constant for bone and brain are not significantly different than muscle and skin (Gabriel et al., 1996b) we expect delays in the same order of magnitude.

Interferential stimulation in (Grossman et al. 2017) was performed mostly at 2 kHz, yet simulations in that work used conductivity values quoted in the literature for 1 Hz. The conductivity values of biological tissues differ between 1 Hz and 1 kHz, but are within one order of magnitude (Gabriel et al., 1996a,1996b; Baumann et al., 1997). We repeated our simulations of inferential stimulation with values reported in the literature for 1 kHz. Figure S4 and Figure S5 show that the results are quite similar to what is in the main text for 1 Hz. The only major change of conductivity in this frequency range is that the scalp conductivity is approximately doubled at 1 kHz (Gabriel et al., 1996b), which led to smaller electric field intensities in the brain.

---

**Figure S1:**
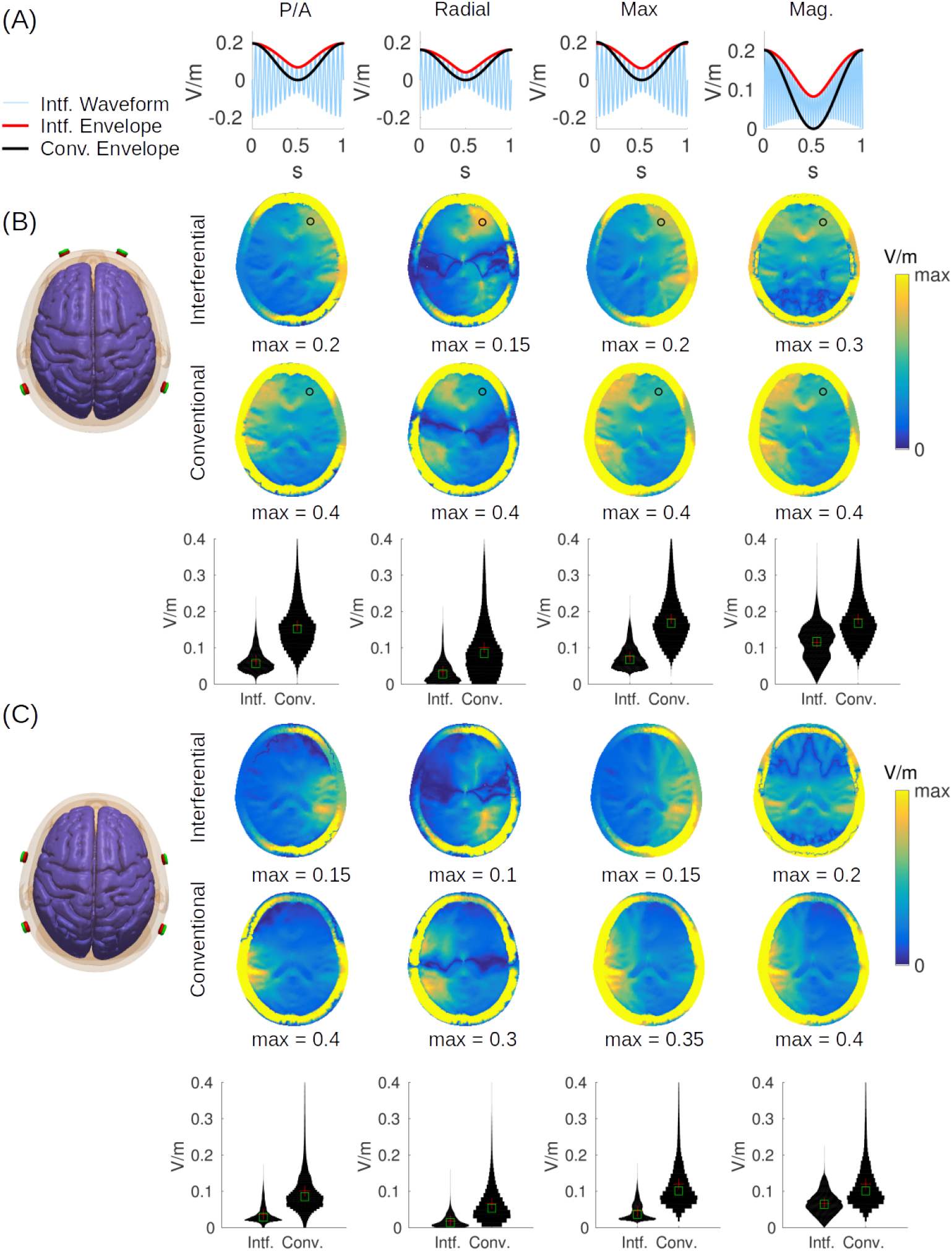
Figure S1: Equivalent of Figure 2, with input currents having ratio of 4/1 between left/right electrode pairs. The total injected current is still kept at 2 mA.

**Figure S2:**
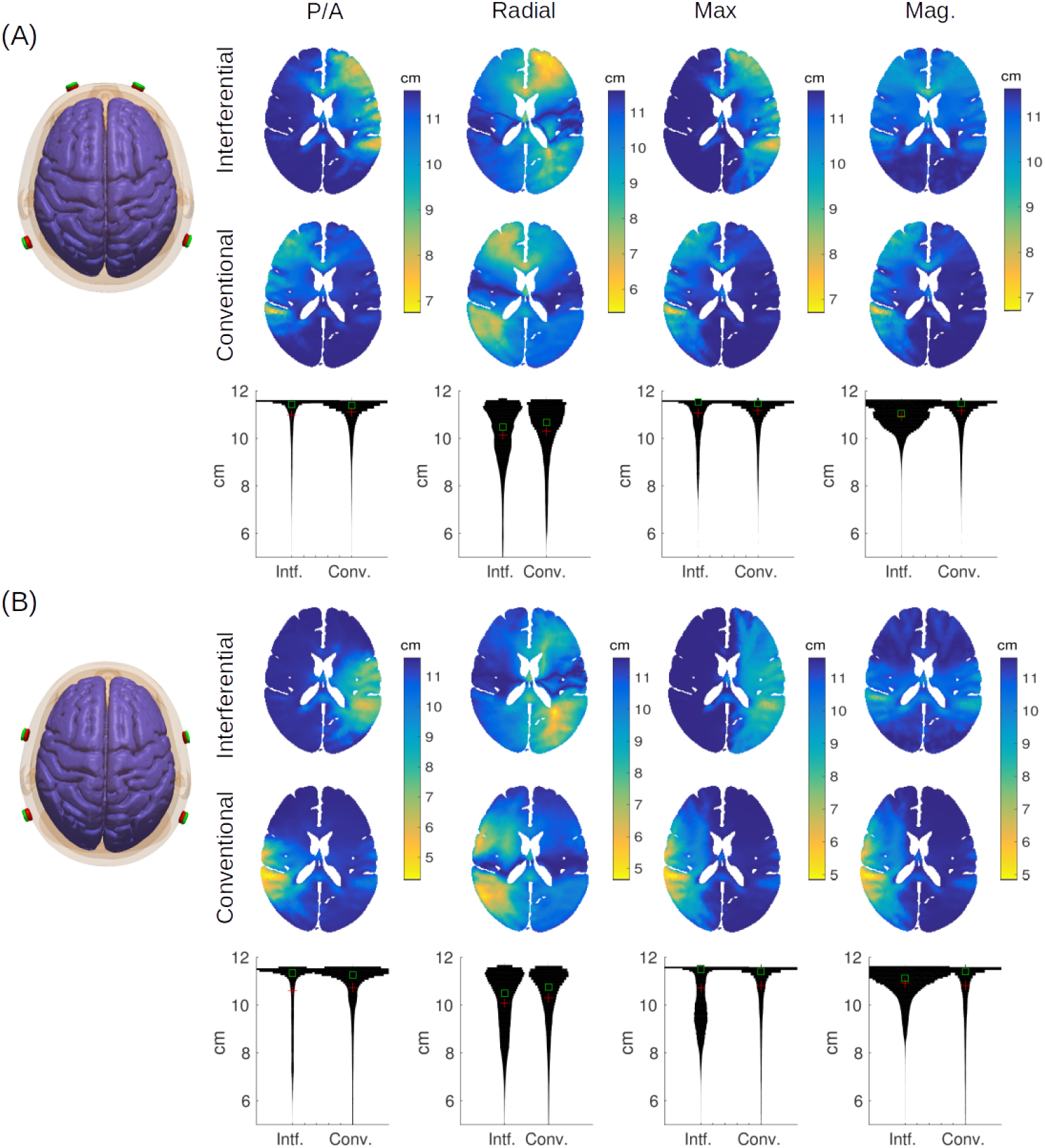
Figure S2: Equivalent of Figure B.4, with input currents having ratio of 4/1 between left/right electrode pairs. The total injected current is still kept at 2 mA.

**Figure S3:**
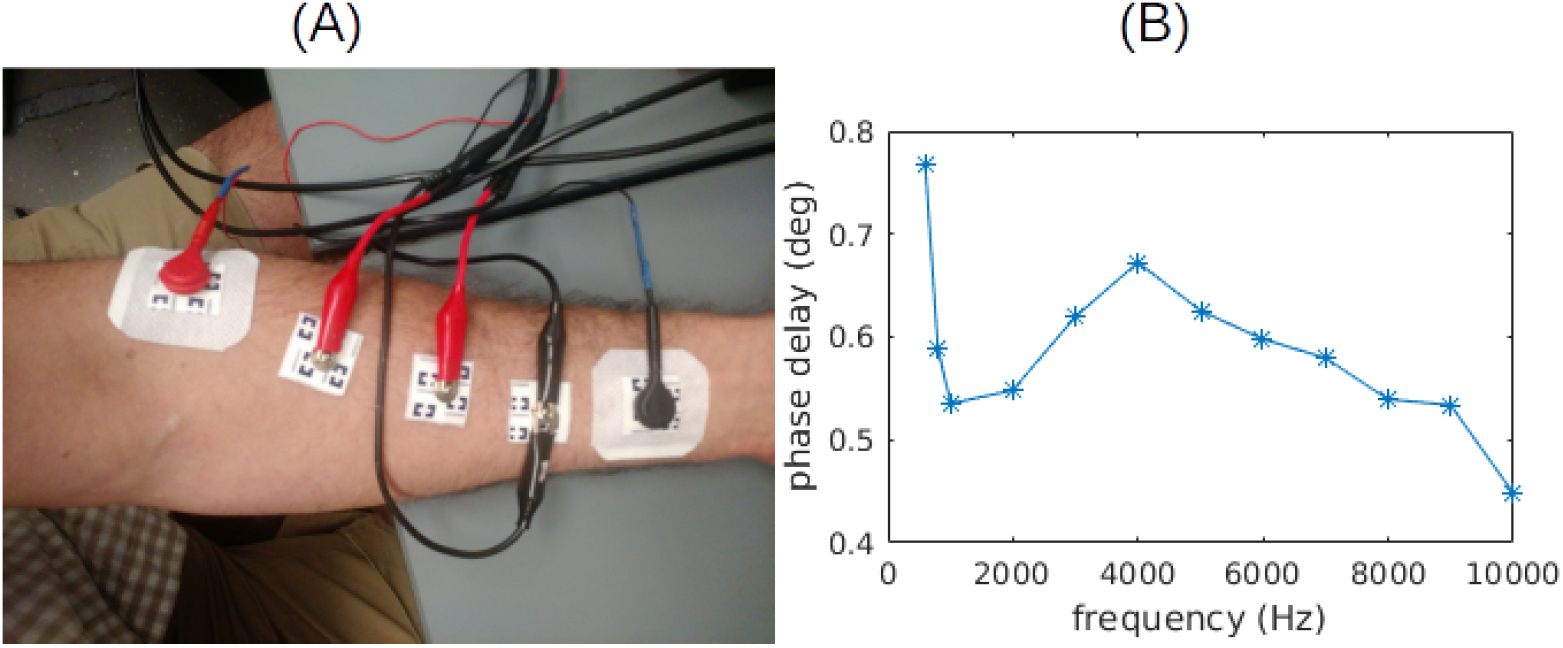
Figure S3: (A) Experimental setup to measure *in vivo* capacitive effects in human tissue. Sinusoidal voltages are recorded at two locations (red alligator clips) relative to a common reference (black alligator clips). Sinusoidal current is applied through distal electrodes (red/black snap electrode connectors) (B) Phase delay between two recorded voltage signals as a function of frequency.

**Figure S4:**
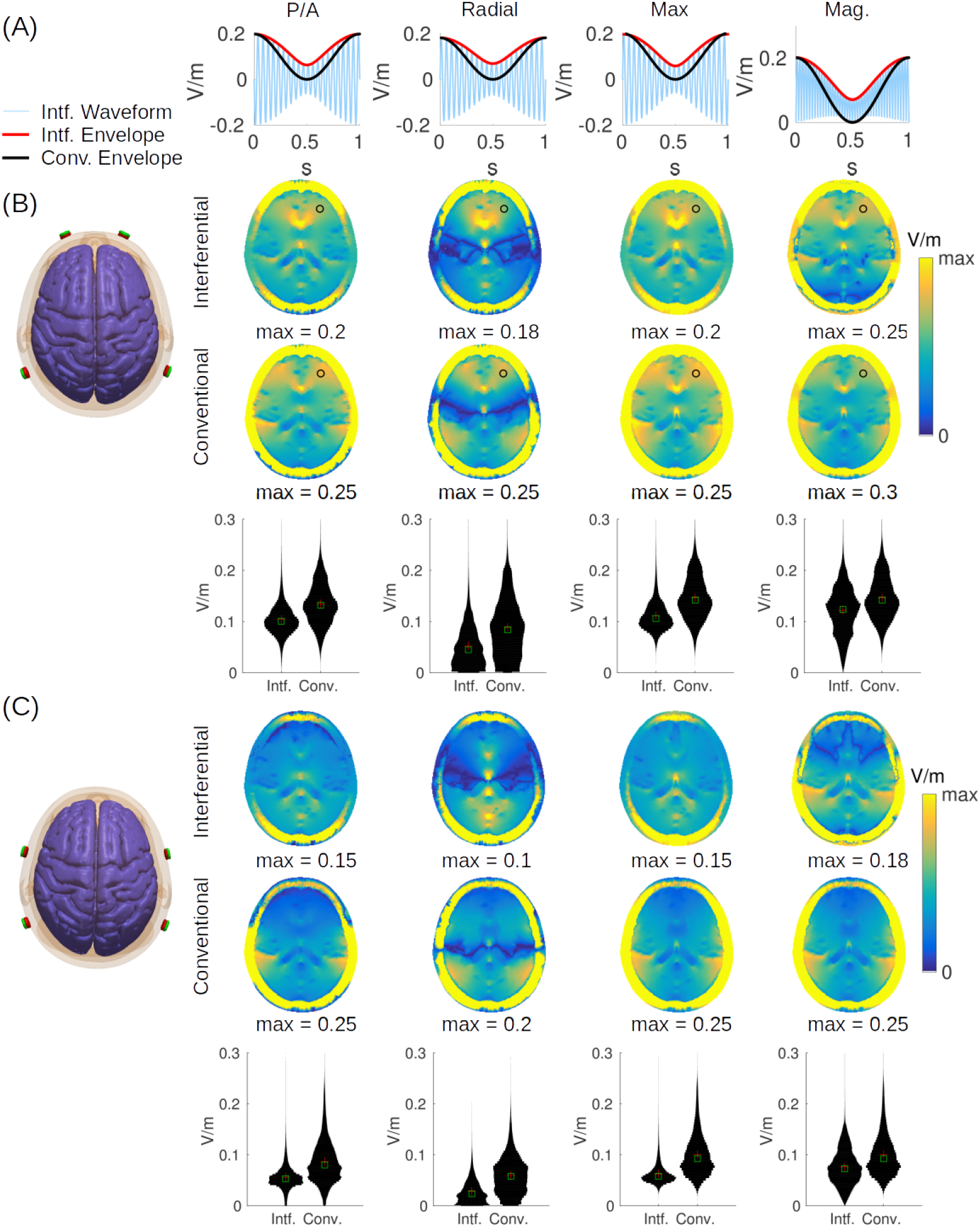
Equivalent of Figure 2, with tissue conductivities for 1 kHz.

**Figure S5:**
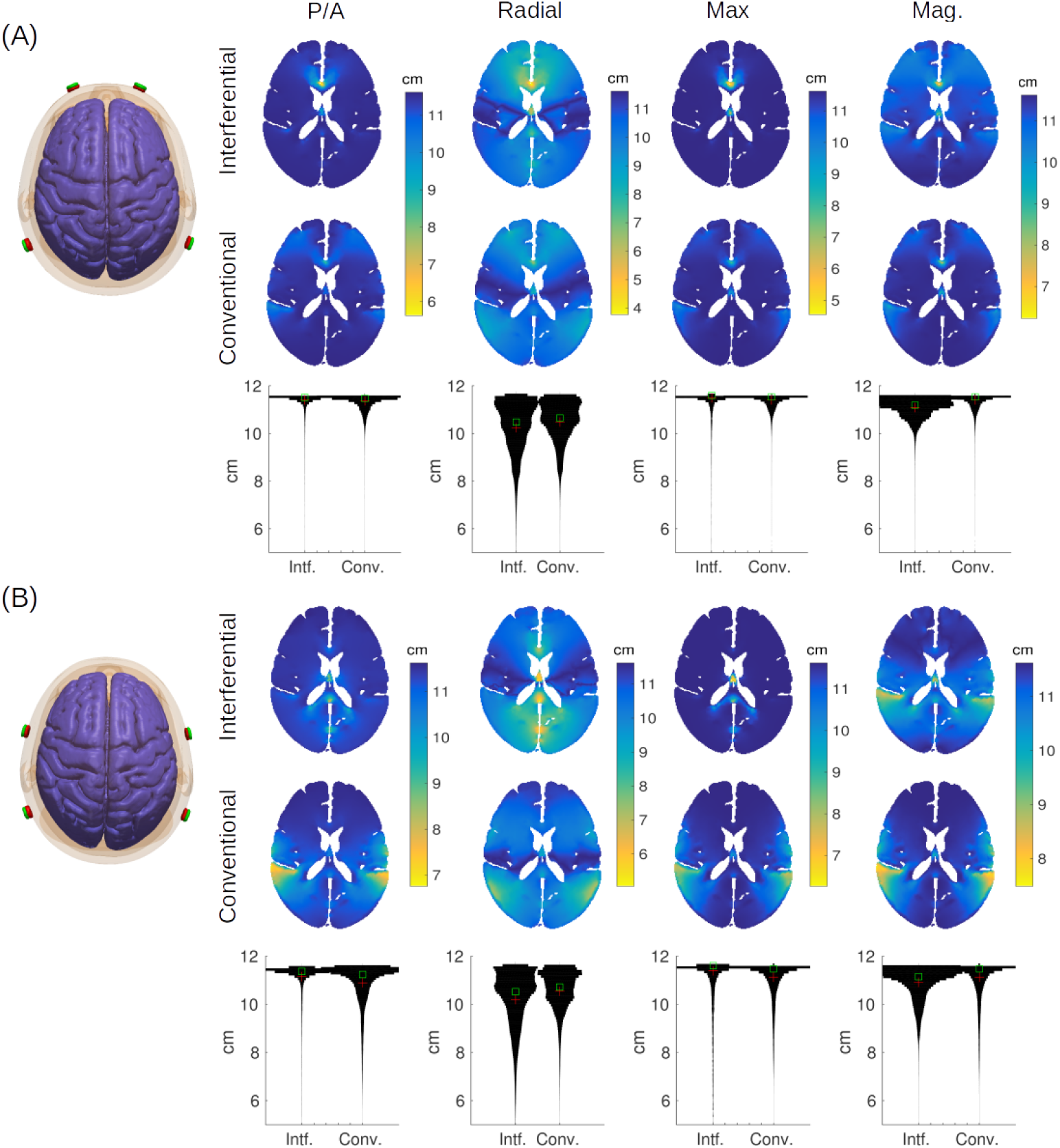
Equivalent of Figure B.4, with tissue conductivities for 1 kHz.

